# Glutathione levels influence chronological life span of *Saccharomyces cerevisiae* in a glucose-dependent manner

**DOI:** 10.1101/217125

**Authors:** Mayra Fabiola Tello-Padilla, Alejandra Yudid Perez-Gonzalez, Melina Canizal-García, Juan Carlos González-Hernández, Christian Cortes-Rojo, Ivanna Karina Olivares-Marin, Luis Alberto Madrigal-Perez

## Abstract

Diet plays a key role in determining the longevity of the organisms since it has been demonstrated that glucose restriction increases lifespan whereas a high-glucose diet decreases it. However, the molecular basis of how diet leads to the aging process is currently unknown. We propose that the quantity of glucose that fuels respiration influences ROS generation and glutathione levels, and both chemical species impact in the aging process. Herein, we provide evidence that mutation of the gene *GSH1* diminishes glutathione levels. Moreover, glutathione levels were higher with 0.5% than in 10% glucose in the *gsh1Δ* and WT strains. Interestingly, the chronological life span (CLS) was lowered in the *gsh1Δ* strain cultured with 10% glucose but not under dietary restriction. The *gsh1Δ* strain also showed an inhibition of the mitochondrial respiration in 0.5 and 10% of glucose but only increased the H_2_O_2_ levels under dietary restriction. These results correlate well with the GSH/GSSG ratio, which showed a decrease in *gsh1Δ* strain cultured with 0.5% glucose. Altogether these data indicate that glutathione has a major role in the function of electron transport chain (ETC) and is essential to maintain life span of *Saccharomyces cerevisiae* in 10% glucose.

## Introduction

Aging is an irreversible process that all the living organisms undergo. It has been postulated that aging leads to the malfunction of multiple cellular processes, which drives to chronic degenerative diseases (Barzilai *et al.*, 2012). Diet is a chief environmental factor that influences the development of chronic degenerative diseases related to aging (Brandhorst *et al.*, 2015; Honma *et al.*, 2012; Prasad *et al.*, 2012; Wei *et al.*, 2017). Accordingly, dietary restriction increases the longevity across a wide range of species and has also been associated with the amelioration of some chronic degenerative diseases (Colman *et al.*, 2009; Colman *et al.*, 2014; Marchal *et al.*, 2012), although it is not entirely clear how the restriction of certain nutrients such as glucose contributes to longevity extension. *Saccharomyces cerevisiae* is a useful model to elucidate the relationships between nutrient load and aging, as the effects of a high-energy diet and dietary restriction can be mimicked in this yeast by evaluating the effects of glucose concentrations of > 1% *w/v* or < 0.5% *w/v*, respectively, on chronological lifespan (CLS), which is the viability of the cell culture in a certain time in a nondividing, quiescent-like condition (Kaeberlein, 2010; Madrigal-Perez *et al.*, 2016; Rockenfeller and Madeo, 2010). High concentrations of glucose decrease CLS by a mechanism related to increased oxidative stress (Barros *et al.*, 2004; Mesquita *et al.*, 2010; Weinberger *et al.*, 2010).

Glutathione (γ-glutamyl-L-cysteinyl-glycine) is the most abundant thiol tripeptide and is involved in numerous cellular processes such as the control of redox environment, iron metabolism, protein stabilization, mitochondrial function, among others (Brasil *et al.*, 2013; Kumar *et al.*, 2011; Penninckx and Elskens, 1993). The levels of glutathione increase under dietary restriction (Rebrin *et al.*, 2003; Sharma *et al.*, 2010; Walsh *et al.*, 2014). Accordingly, deletion of *SCH9*, *TOR1*, and *RAS2* nutrient-sensing genes exhibits a dietary restriction phenotype and maintain high levels of reduced glutathione when are exposed to stress, besides displaying enhanced survival to the anti-cancer agent cisplatin by using glutathione to detoxify this drug (Mariani *et al.*, 2014). Overall, these findings suggest that glutathione levels are regulated by glucose or other nutrients and may have a fundamental role during cellular stress.

Glutathione is synthesized in a two-step process catalyzed by the proteins encoded by the genes *GSH1* and *GSH2* (Tang *et al.*, 2015). The mutation in *S. cerevisiae* of the gene *GSH1* (*gsh1Δ*) dramatically decreases glutathione levels from approximately 4 mM to 0.03 mM and increases yeast sensitivity to H_2_O_2_ (Hatem *et al.*, 2014). The *gsh1∆* mutation also reverted the resistance of *S. cerevisiae* to the anti-cancer compound cisplatin under dietary restriction (Mariani *et al.*, 2014).

Considering that dietary restriction increases glutathione levels and extends longevity and that glutathione optimizes the mitochondrial function, the aim of this work was to elucidate whether longevity extension by dietary restriction is mediated by an improvement of mitochondrial function via an enhancement of both the redox status and the concentration of glutathione.

## Material and methods

### Strains

The BY4741 wild type strain of *S. cerevisiae* (*MATa; his3Δ1; leu2Δ0; met15Δ0; ura3Δ0;* WT) and its *gsh1*Δ mutant *(MATa; his3Δ1; leu2Δ0; met15Δ0; ura3Δ0; YJL101c:: KanMX4; gsh1Δ*) were acquired from EUROSCARF (Frankfurt, Germany). The strains were maintained in yeast extract-peptone-dextrose (YPD) medium (1% yeast extract, 2% casein peptone and 2% glucose). Medium for the mutant strain was supplemented with 150 μg/mL of G-418 disulfate salt (Sigma-Aldrich, St. Louis, MO, USA).

### Chronological life span assay

The CLS was performed according to the protocol described by Murakami *et al.* (2008) as follow. WT and *gsh1Δ* cells were cultured from frozen stocks by plating onto YPD agar, consisting of 1% yeast extract (Sigma-Aldrich), 2% casein peptone (Sigma-Aldrich), 2% bacteriological agar (Affymetrix, Santa Clara, CA, USA) and supplemented with 2% of glucose. The plates were incubated at 30ºC for 48 h or until single colonies appear. Then, three singles colonies were picked, and cultured overnight in 2 mL YPD medium supplemented with 2% glucose. To induce aging in the cultures, 300 µL of the fresh overnight culture were placed into 3 mL YPD (0.5, 2 or 10% glucose) medium in a 15 mL test tube to keep 1:5 liquid-air ratios. Cultures were maintained at 30 ºC with constant agitation at 180 rpm throughout the experiment. After incubation for 2 days, 500 µL aliquots were taken from aging cultures and inoculated into 5 mL YPD medium supplemented with 2% glucose in a 25 mL Erlenmeyer flask. Cultures were incubated at 30 ºC for 12 h in a shaking incubator (MaxQ 4000, Thermo Scientific, MA, USA) at 180 rpm. The absorbance of the cultures at 600 nm was measured every hour during a period of 12 h using a UNICO spectrophotometer (UNICO, NJ, USA). The growth kinetics procedure was repeated every two days. To determine the survival percentage, the doubling time (*td*) was calculated by adjusting the growth curves with the exponential growth equation using the software GraphPad Prism 6.00 (GraphPad Software, La Jolla, CA, USA). Additionally, the time shift (*Δtn*) was obtained by calculating the natural logarithm of the optic density at 600 nm (OD_600_) obtained at the exponential phase of the growth curve and then interpolating at OD_600_ = 0.5 using linear regression. Finally, the survival percentage (*Sn*) was calculated according to the following equation:

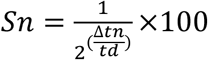

On the other hand, the area under curve (AUC) survival was computed from the data of survival percentage vs. time using the trapezoid rule in the GraphPad Prism 6.00.

### Determination of in situ mitochondrial respiration

The oxygen consumption was measured polarographically at 28 ºC using a Clark-type oxygen electrode (YSI 5300, Yellow Springs, OH, USA). *S. cerevisiae* cultures were harvested either at mid-log phase (D.O. _600_ ~ 0.5) or after six days of inoculation (chronologically aged yeast culture) at 5000 x *g* for 5 minutes at 28 ºC. Then, 125 mg of cells were resuspended in 5 mL of buffer 10 mM 2-(N-morpholino) ethane sulfonic acid (MES-TEA; Sigma-Aldrich), adjusted to pH 6.0 with triethanolamine (Sigma-Aldrich) in a closed chamber with constant stirring. The basal respiration was determined using 10 mM glucose as substrate. The maximal rate of respiration was evaluated by adding 10 µM of the uncoupler carbonyl cyanide 3-chlorophenylhydrazone (CCCP). Non-mitochondrial oxygen consumption was discriminated by adding 10 µM Antimycin A at the end of each determination. Oxygen consumption was expressed as nanoatoms O_2_ per minute per milligram of cells.

### Quantification of H_2_O_2_ release

Superoxide anion produced in the mitochondrial matrix is rapidly dismutated by the manganese-dependent superoxide dismutase (Sod2p) to hydrogen peroxide (H_2_O_2_), which diffuses through mitochondrial and plasmatic membranes to the surrounding medium (Dikalov and Harrison, 2014). Thereby, the extracellular H_2_O_2_ release was used as an indicator of ROS levels and was determined by the Amplex red hydrogen peroxide assay kit (Invitrogen, Waltham, MA, USA) according to the manufacturer’s instructions. Briefly, overnight *S. cerevisiae* cultures were harvested at 5000 x *g* for 5 minutes at 28 ºC. The cells were washed three times with deionized water and resuspended in 2 mL of assay buffer containing 20 mM Tris-HCl, 0.5 mM EDTA and 2% ethanol at pH 7. The cells were placed in 96 well black plates at a density of 3 × 10^6^ cells/well. Then, 50 µL of work solution (100 µM Amplex red and 10 U/mL horseradish peroxidase) was added to each well and incubated for 30 min at 30ºC with constant agitation in a horizontal shaker and protected from light. Finally, the basal release of H_2_O_2_ was measured at an excitation wavelength of 563 nm and an emission wavelength of 587 nm with a microplate reader (Varioskan Flash Multimode Reader, Thermo-Scientific).

### Quantification of glutathione

Total glutathione, which comprises the sum of the concentrations of reduced glutathione (GSH) plus oxidized glutathione (GSSG), was quantified by measuring the reduction of 5,5 ′-dithio-bis(2-nitrobenzoic acid) (DTNB) (Rahman *et al.*, 2006; Tietze, 1969). 3 mL of exponential phase cultures (OD_600_ ∼ 0.5) or chronologically aged cultures (six days after inoculation) were harvested at 5000 x *g* for 2 min. Cells were normalized to an OD_600_ ∼ 0.5, washed twice with 2 mL PBS, resuspended in 25 µL of PBS and homogenized by vortexing with glass beads. Then, the cytosolic extracts were deproteinized with 5-sulfosalicylic acid 5% (Sigma-Aldrich) at −20ºC for 30 min. Glutathione determination reaction was prepared using 700 μL buffer KPE (0.1 M phosphate potassium, 0.001 M EDTA), 100 μL cell extract, 60 µL glutathione reductase solution (6 U/mL; Sigma-Aldrich), 60 µL of 0.66 mg/mL DTNB solution and 60 µL of 0.66 mg/mL β-NADPH solution (Sigma-Aldrich). Finally, the samples were read at 412 nm using a microplate reader (Varioskan Flash Multimode Reader, Thermo-Scientific). For GSSG determination, the cell extracts were treated with 4-vinyl pyridine (Sigma-Aldrich) for GSH derivatization, and the excess of 4-vinyl pyridine was neutralized with triethanolamine. Then, GSSG was quantified by reducing it with glutathione reductase as described above.

### Statistical analyses

Two-tailed unpaired Student *t-*test was applied to test for differences in CLS, H_2_O_2_ release, total glutathione concentration and rate of oxygen consumption. For all analyses, at least 3 independent experiments were performed. Statistical analyses were computed using GraphPad Prism 6.00 (GraphPad Software, La Jolla California, USA).

## Results

### Effect of glucose concentration on H_2_0_2_ levels and mitochondrial function in the exponential growth phase and its impact over CLS

As shown in the Fig. 1a, 10% glucose decreases the yeast CLS severely, while dietary restriction, which was mimicked by adding 0.5% glucose to the culture media, increases in a notable way the CLS, as reflected by a full preservation of cell viability all over the experiment. Antimycin A-sensitive respiration and H_2_O_2_ release were measured to assess if the effects of excessive glucose were related to differences in the exponential growth phase (i.e. before the beginning of chronological aging) in mitochondrial function and ROS production, respectively. It can be observed a 60% increase of H_2_O_2_ levels in the yeast growing with 10% glucose in comparison to yeast with 0.5% glucose (Fig. 1b), as well a severe repression of respiration in both state 4 (i.e. basal respiration) and uncoupled (U) state (i.e. maximal respiration rate with CCCP). Besides, respiration in cells under dietary restriction was almost entirely uncoupled as the increase of oxygen consumption elicited by CCCP was negligible (Figs. 1c and d). Therefore, these results suggest that excessive concentrations of glucose during the exponential growth set early defects in mitochondrial function that may be related to exacerbated ROS production and/or antioxidant depletion that contribute to the phenotype of impaired CLS.

**Figure 1.**
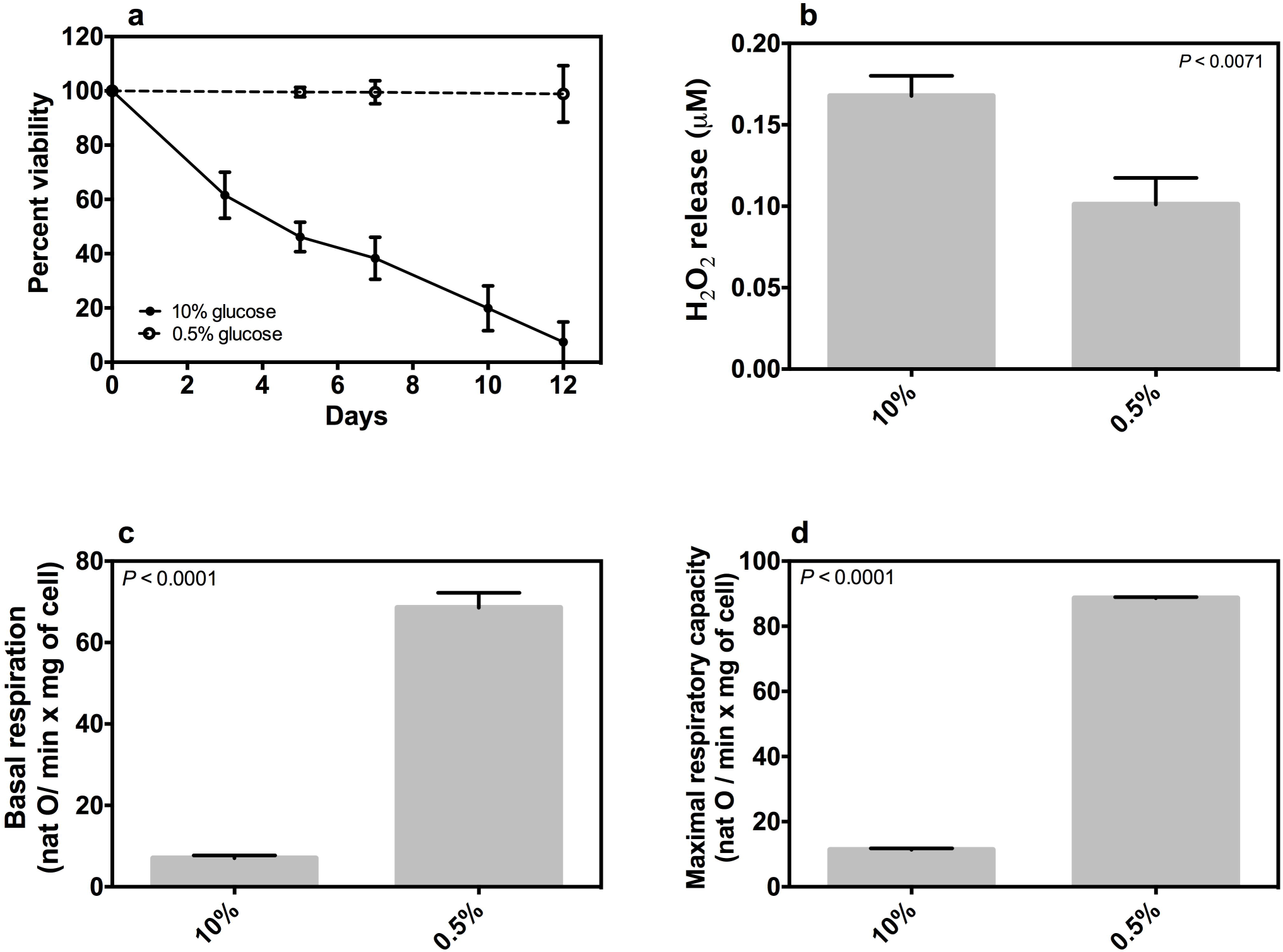
Effect of glucose concentration on the chronological life span and cellular bioenergetics of *S. cerevisiae*. **a)** The chronological life span of *S. cerevisiae* was measured in SC minimal media supplemented with 0.5 or 10% of glucose; **b)** the H_2_O_2_ release was measured by Amplex Red kit in *S. cerevisiae* cells grown in YPD medium supplemented with 0.5 or 10% of glucose; **c-e)** the oxygen consumption was measured at basal state and maximal respiratory capacity in *S. cerevisiae* cells grown in YPD medium supplemented with 0.5 or 10% of glucose. The results represent mean values ± SEM from 3-5 independent experiments, which includes mean values of 3 technical repetitions. Statistical analyses were performed using two-tailed unpaired Student *t-*test.

### Effect of GSH1 mutation on glutathione levels at the exponential growth phase

The levels of glutathione in the WT cells and the *gsh1Δ* mutant were compared at three concentrations of glucose in order to explore if the levels of this antioxidant at the exponential growth phase are influenced by the concentration of glucose. As expected, total glutathione content decreased ~50% in the *gsh1Δ* strain in comparison to WT cells at each level of glucose tested (Fig. 2). It is also important to mention that the *gsh1Δ* strain is glutathione auxotroph and incapable of growing in minimal media. For this reason, we have used YPD medium. It is pertinent to note that in the mutant strain the glutathione was not completely depleted and this may be due to the YPD media glutathione content. 10% glucose decreased glutathione concentration in the WT cells at similar levels to that observed in the *gsh1Δ* mutant at 0.5% and 2% glucose. Regarding to the *gsh1Δ* mutant, a ~50% decrease in comparison to WT was observed only a 10% of glucose. These data suggest that compromised CLS may be the result of impaired mitochondrial function and excessive ROS production due to severe diminution of glutathione at the exponential growth phase.

**Figure 2.**
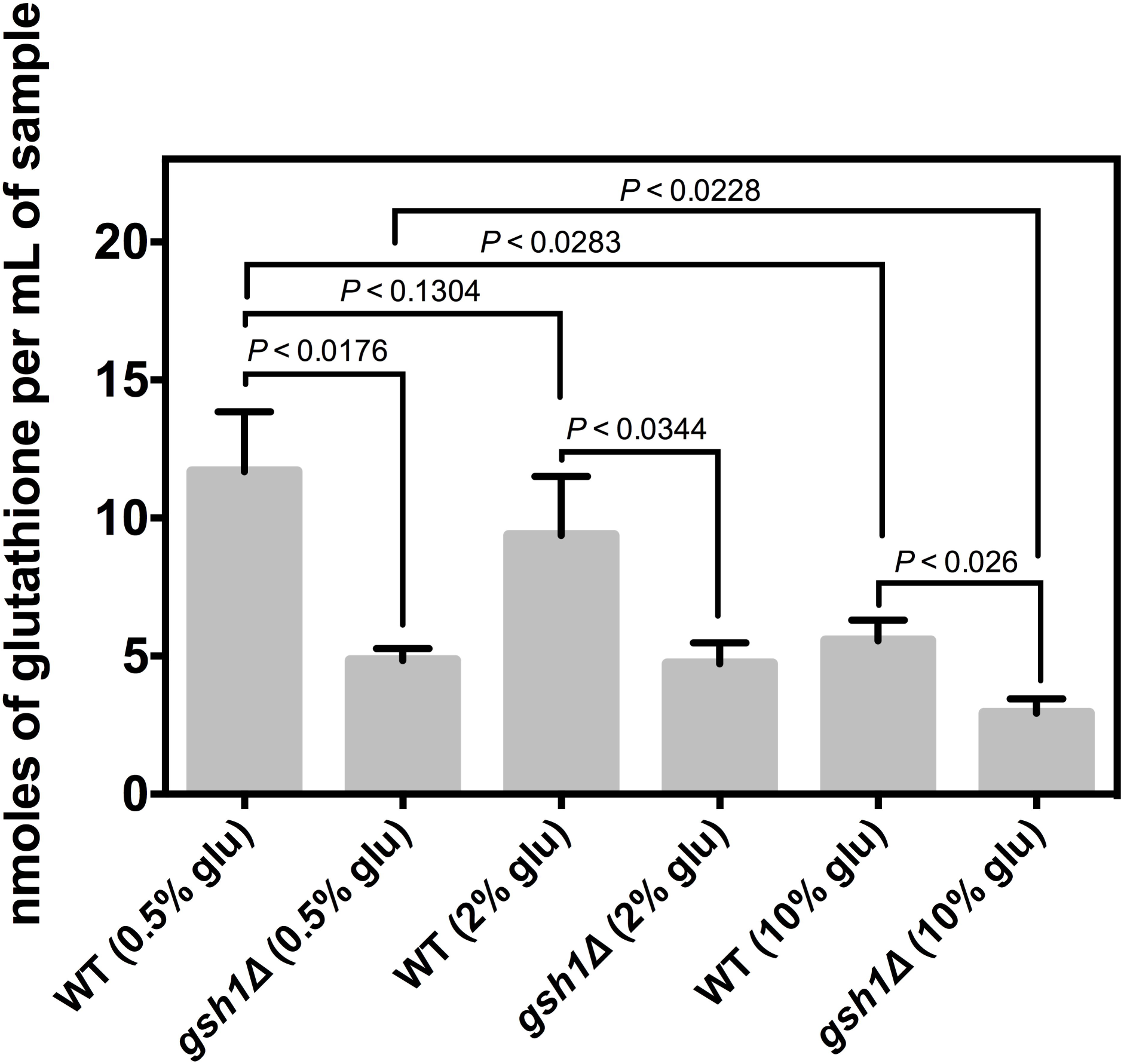
Effect of glucose concentration and mutation of the gene *GSH1* in the total glutathione of *S. cerevisiae.* Total glutathione was quantified by recycling assay in the WT and *gsh1Δ* strains grown in YPD media supplemented with 0.5, 2 or 10% of glucose. Statistical analyses were performed using two-tailed unpaired Student *t-*test.

### Influence of the GSH1 mutation on both H_2_O_2_ levels and CLS

The effect of the *GSH1* mutation on H_2_O_2_ levels was tested under dietary restriction and both standard and high-glucose levels. As seen in the Fig 3, ROS levels remained high in the *gsh1Δ* mutant irrespectively of the glucose concentration tested. In contrast, a ~40% decrease in ROS levels was observed in the WT strain under dietary restriction when compared to the *gsh1Δ* mutant. These data suggest that intact glutathione synthesis is required for the decrement of ROS levels elicited by dietary restriction.

**Figure 3.**
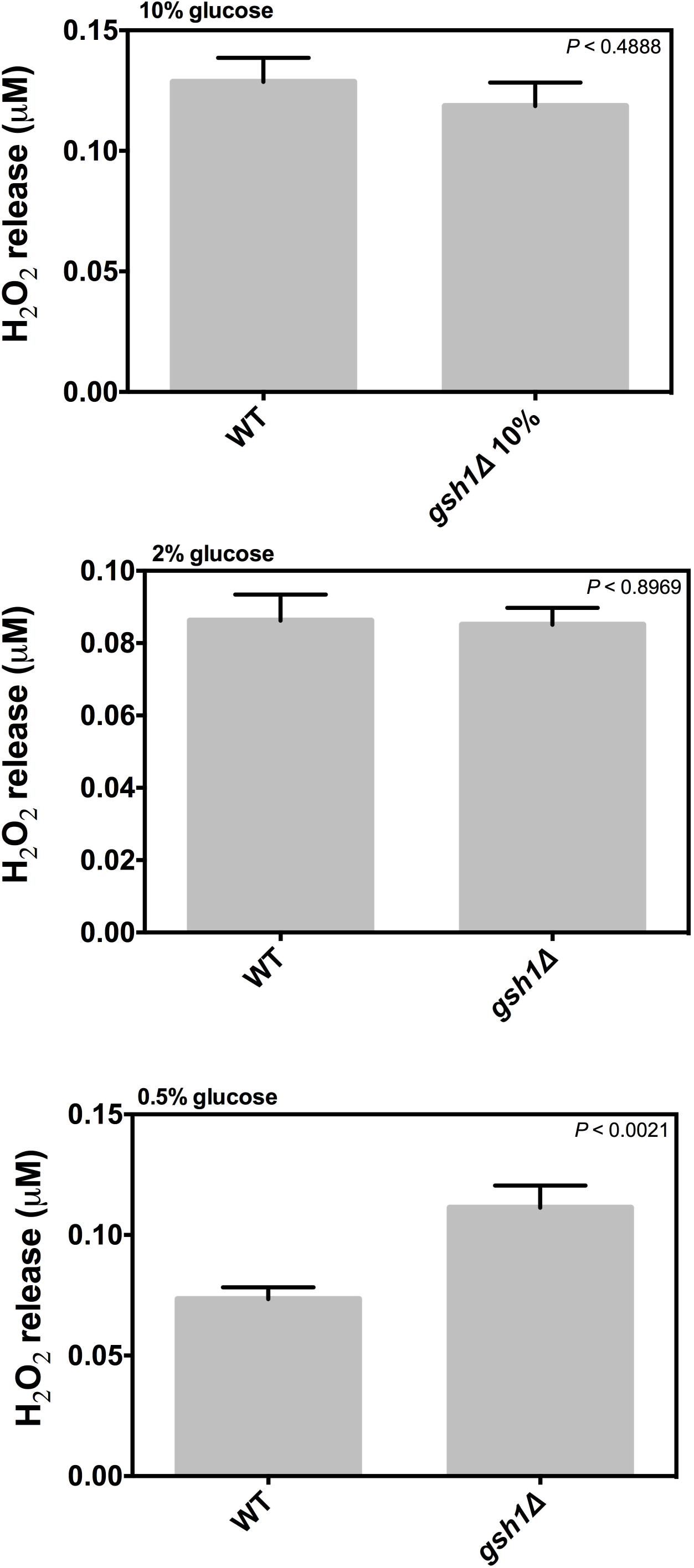
Influence of glucose concentration and mutation of the gene *GSH1* on the H_2_O_2_ release of *S. cerevisiae.* The Amplex Red kit was used in the WT and *gsh1Δ* strains grown in YPD medium supplemented with 0.5, 2 or 10% of glucose to measure the H_2_O_2_ release. Results represent mean values ± SEM from 5 independent experiments. Statistical analyses were performed using two-tailed unpaired Student *t-*test.

The Fig 4 shows the CLS of both the WT and *gsh1Δ* strains. At high-glucose level (i.e. 10%) a drastic decrease of CLS was detected in the *gsh1Δ* mutant in comparison to the WT cells. However, no differences in CLS were observed among the WT cells and the *gsh1Δ* mutant at the standard-glucose level and under dietary restriction. On the other hand, CLS was tested in the presence of 9.5 % sorbitol, a non-metabolizable carbohydrate plus 0.5% glucose to discard that decreased CLS in the *gsh1Δ* mutant was due to an osmotic effect instead of a result of nutrient overload. As expected, this approach caused a longer CLS in the *gsh1Δ* mutant the exposure to 10% glucose (Fig. 5). This result indicates that decreased CLS in the *gsh1Δ* mutant is not attributable to an osmotic stress elicited by the inability of this strain to synthesize glutathione.

**Figure 4.**
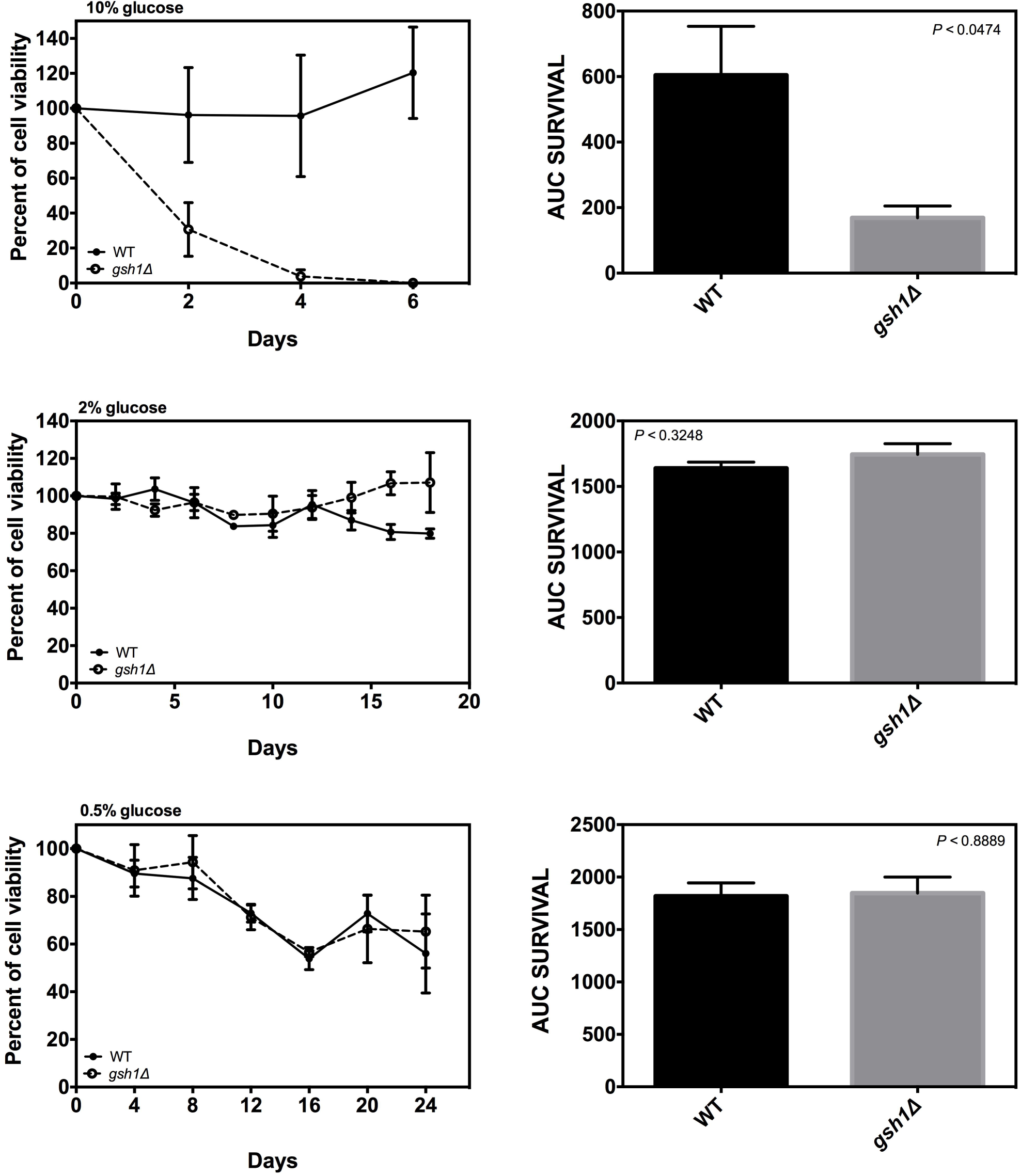
Effect of glucose concentration and mutation of the gene *GSH1* in the chronological life span of *S. cerevisiae.* The chronological life span of WT and *gsh1Δ* strains was measured in YPD medium supplemented with 0.5, 2 or 10% of glucose. The area under curve (AUC) survival was computed from the data of survival percentage vs. time using the trapezoid rule in the GraphPad Prism 6.00 for Macintosh. The results represent mean values ± SEM from 4 independent experiments, which includes mean values of 3 technical repetitions. Statistical analyses were performed using two-tailed unpaired Student *t-*test.

**Figure 5.**
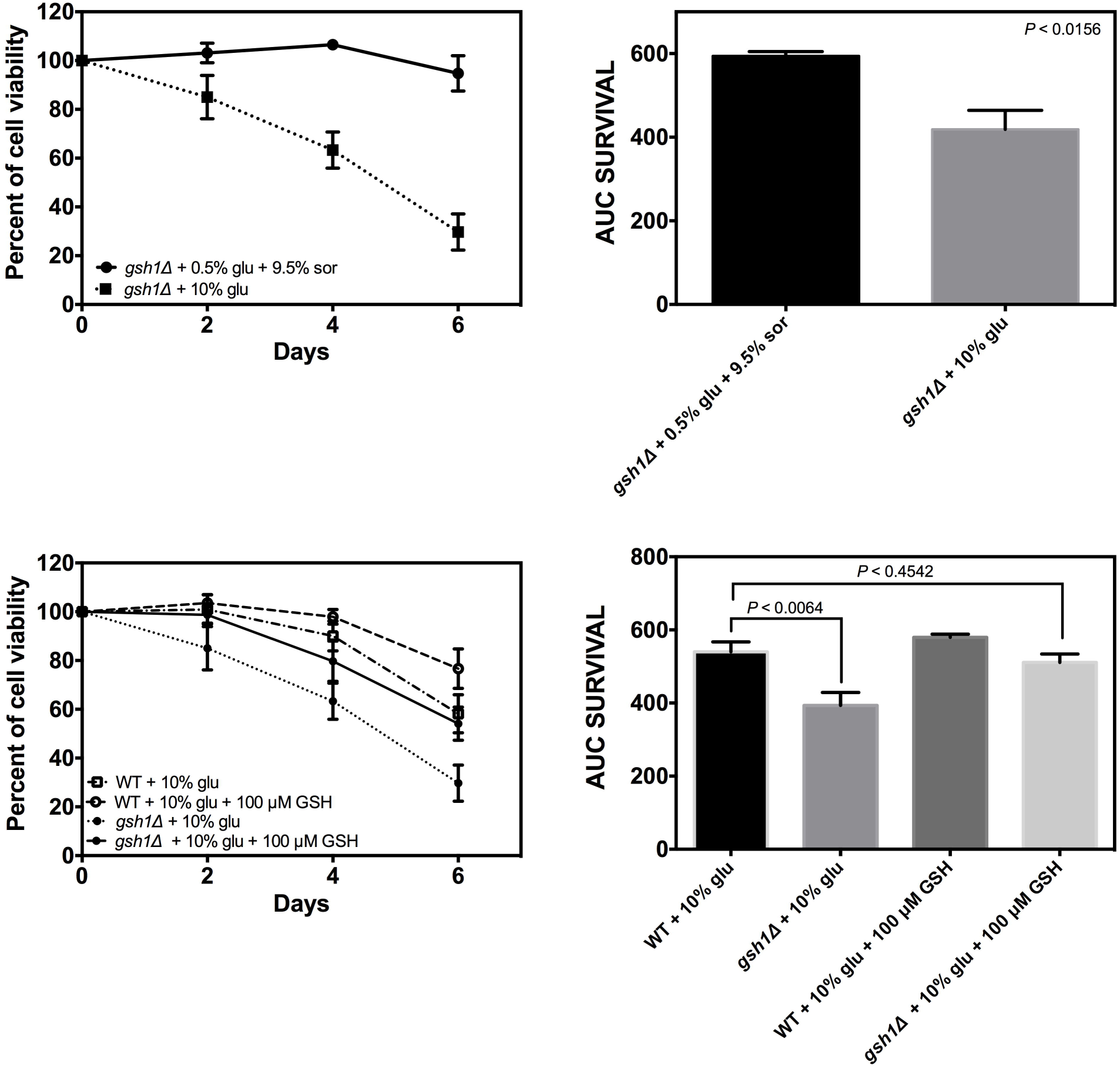
Influence of sorbitol, glutathione supplementation, and mutation of the gene *GSH1* in the chronological life span of *S. cerevisiae.* The chronological life span of WT and *gsh1Δ* strains was measured in YPD medium supplemented with 0.5% glucose (glu) plus 9.5% sorbitol (sor) and 10% of glucose supplemented with 100 µM glutathione (GSH). The area under curve (AUC) survival was computed from the data of survival percentage vs. time using the trapezoid rule in the GraphPad Prism 6.00 for Macintosh. The results represent mean values ± SEM from 6 independent experiments. Statistical analyses were performed using two-tailed unpaired Student *t-*test.

Supplementation of the culture media with 100 µM glutathione was carried out to assess what extent the diminution of the CLS observed in the *gsh1Δ* mutant at 10% glucose was the consequence of decreased glutathione content. Glutathione addition increased the CLS of the *gsh1Δ* mutant at the same level than the WT strain at 10% glucose (Fig. 5).

### Influence of GSH1 mutation over the mitochondrial function of yeast during CLS

The *in situ* mitochondrial respiration of *S. cerevisiae* was measured to determine if the decrease of the CLS of *gsh1Δ* mutant at 10% glucose was the result of impaired mitochondrial function.

In the exponential phase cultures, a severe decrease in both state 4 and U states was found in the *gsh1Δ* strain at 0.5 and 10% glucose in comparison to the WT cells (Fig. 6). Indeed, oxygen release was observed at 10% glucose in the *gsh1Δ* strain, which is indicative of exacerbated ROS production. In contrast, no differences were seen in the respiratory states of both WT and *gsh1Δ* cells at 2% glucose (Fig. 6). Unexpectedly, it was observed that sorbitol augmented several folds the respiratory rates in both state 4 and state U when compared to the *gsh1Δ* mutant growth only with 0.5% glucose, although respiration in U state did not reach the levels observed in the WT cells growth also with 0.5% glucose and 9.5% sorbitol (Fig. 6). It is also remarkable that this treatment elicited a more uncoupled respiration in the *gsh1Δ* mutant than in the WT cells as the ratio state U/state 4 was ~2 and ~3, respectively.

**Figure 6.**
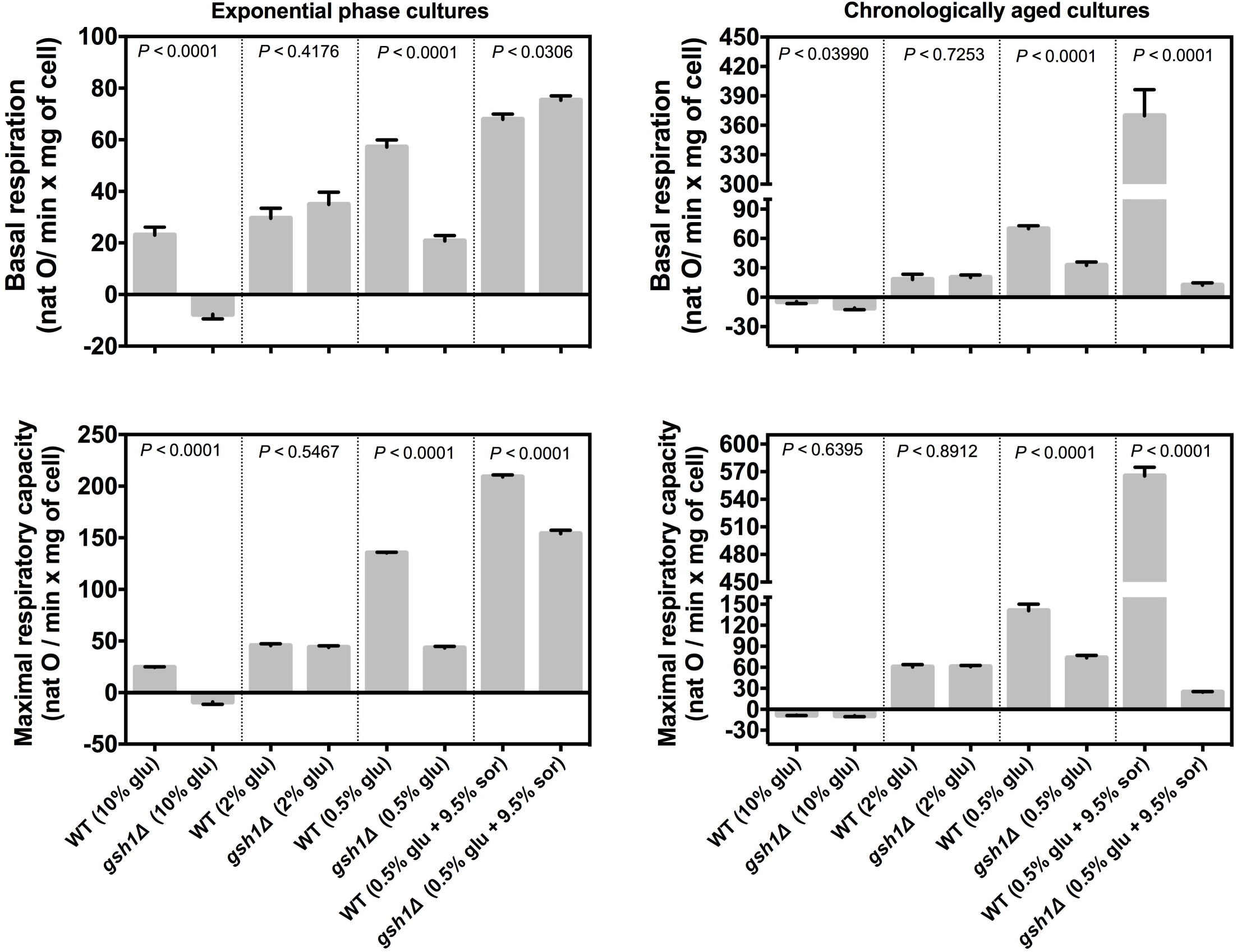
Influence of glucose concentration, sorbitol, and mutation of the gene *GSH1* in the mitochondrial respiration of *S. cerevisiae.* The oxygen consumption was measured at basal state and maximal respiratory capacity in *S. cerevisiae* in exponential phase and chronologically aged grown in YPD medium supplemented with 0.5, 2 or 10% of glucose (glu) and 0.5% glucose plus 9.5% sorbitol (sor). The results represent mean values ± SEM from 4 independent experiments, which includes mean values of 3 technical repetitions. Statistical analyses were performed using two-tailed unpaired Student *t-*test.

Regarding to mitochondrial function during chronological aging, both WT and *gsh1Δ* cells exhibited a full inhibition of respiration in any respiratory state with 10% glucose, being observed instead oxygen release to the medium (Fig. 6). Also, no differences were detected in any respiratory state at 2% glucose; however, a 50% decrease in respiration was observed in the *gsh1Δ* mutant with respect to WT cells under dietary restriction. On the other hand, divergent results were obtained when sorbitol was added during dietary restriction, since the respiration of the WT strain increased several times at levels even higher than that observed in the exponential phase, while in the *gsh1Δ* mutant, sorbitol severely depressed respiration at the lower levels among all the experimental conditions (Fig. 6). Overall, these results suggest that either during the exponential phase or the chronological aging, *GSH1* is needed to achieve an optimal mitochondrial function during dietary restriction, but it has not influence at higher concentrations of glucose.

### Effects of GSH1 mutation on oxidative stress during CLS

To explore whether enhanced oxidative stress due to *GSH1* mutation is involved in the shortening of yeast CLS in high glucose we measured the GSH/GSSG ratio. As expected, oxidative stress augmented drastically in WT cells by the addition of 10% glucose in comparison to dietary restriction, as the GSH/GSSG ratio was more than twice lower at 10% glucose than at 0.5% (Fig. 7). In the *gsh1Δ* mutant, GSH/GSSG ratio was severely diminished under dietary restriction in comparison to the WT strain, but non-statistically significant differences were observed among these strains at 10% glucose. These data suggest that intact glutathione synthesis is required to keep oxidative stress at minimum only during calorie restriction.

**Figure 7.**
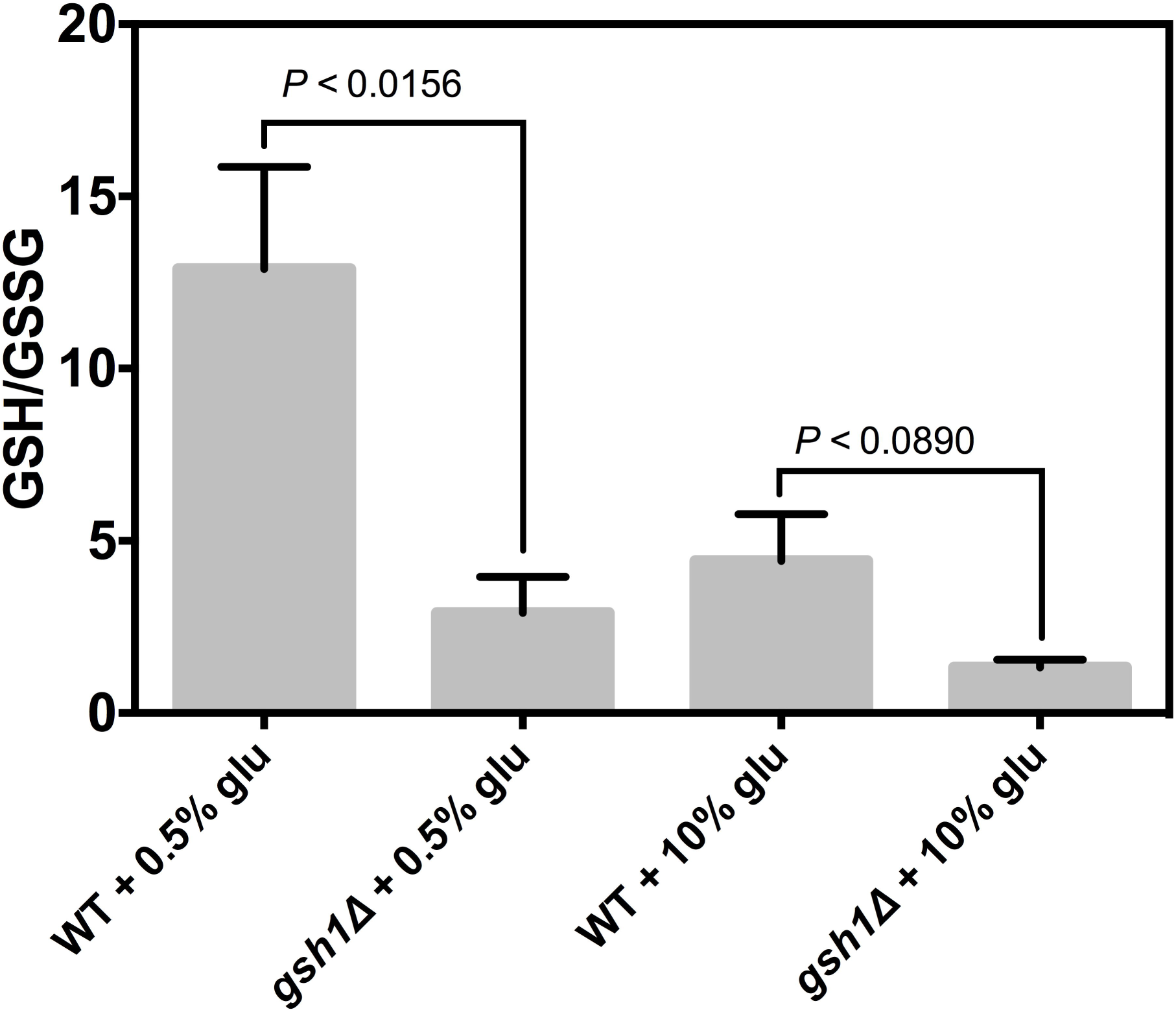
Effect of glucose concentration and mutation of the gene *GSH1* in the GSH/GSSG ratio of *S. cerevisiae.* GSH/GSSG ratio was quantified by recycling assay in the WT and *gsh1Δ* strains grown in YPD media supplemented with 0.5 or 10% of glucose. Statistical analyses were performed using two-tailed unpaired Student *t-*test.

## Discussion

Delaying of the aging process is critical to counteract the onset of age-related diseases and preserve health span (Barzilai *et al.*, 2012). However, the molecular basis of the aging process is currently unknown. Restriction of certain nutrients such as glucose and supplementation with resveratrol improve the longevity of a wide range of species and have also been related with the amelioration of some chronic degenerative diseases (Huffman *et al.*, 2016). Resveratrol and dietary restriction affect the mitochondrial function and the antioxidant systems, which directly impact ROS generation (Barros *et al.*, 2004; Madrigal-Perez *et al.*, 2016; Plauth *et al.*, 2016). Nonetheless, it is unclear whether the molecular mechanism by which dietary restriction counteract oxidative stress also modulates mitochondrial function and the aging process. Thereby, the aim of this study was to determine whether the amount of glucose in the culture medium influences ROS generation and glutathione levels, and its relationship with the aging process.

The mutant *gsh1Δ* of *S. cerevisiae* exhibited diminished glutathione levels independently of the concentration of glucose (Fig. 2), while under dietary restriction (i.e. 0.5% glucose), displayed increased H_2_O_2_ levels (Fig. 3), impaired mitochondrial function either during the exponential growth or during chronological aging (Fig. 6) and enhanced oxidative stress with respect to WT cells (Fig. 7). Nevertheless, its chronological longevity was severely affected only in the presence of 10% glucose (Fig. 4). On the contrary, the effects of glucose on WT cells show a dependence on glucose concentration as seen with glutathione concentrations (Fig. 2), H_2_O_2_ levels (Fig. 3), mitochondrial function (Fig. 6) and oxidative stress (Fig. 7). It was discarded that the deleterious effects of 10% glucose over these parameters were due to an osmotic effect, by supplementing 9.5% sorbitol in the culture with 0.5 glucose (Figs. 5 and 6). However, the most surprising finding from these data is that sorbitol enhanced several times the respiration in the WT cells during chronological aging and, at lower extent, in the logarithmic phase, although in the *gsh1Δ* mutant, sorbitol decreased respiration at any respiratory state (Fig. 6) without affecting CLS (Fig. 5). Osmotic stress activates a myriad of responses via the high-osmolarity glycerol (HOG) pathway (Brewster *et al.*, 1993), among which, augmented catalase T activity is one of them (Schüller *et al.*, 1994). Besides, it has been described that high osmolarity by NaCl increases glutathione synthesis (Jamnik *et al.*, 2006). Assuming that osmotic stress by sorbitol also induces glutathione synthesis, it can be proposed that enhanced respiration was due to enhanced protection against the deleterious effects of aging and glucose on the ETC conferred by the synergic action of the antioxidant responses mounted by both dietary restriction and osmotic stress. In this regard, ROS like H_2_O_2_ decreases the activity of the complex I in mammalian mitochondria via glutathionylation of free cysteine residues as the concentration of oxidized glutathione increases (Taylor *et al.*, 2003). Although *S. cerevisiae* does not contain a canonical complex I, it has been demonstrated that complexes II and III from yeast mitochondria are inhibited by H_2_O_2_ and that this effect can be reverted by a thiol reductant (Cortes-Rojo *et al.*, 2007). Thus, the discrete effect of sorbitol on the respiration of the *gsh1Δ* mutant may be related to that this strain cannot mount a response dependent on glutathione, which might drive to a higher inhibition of the ETC. However, the residual respiratory activity seen in this mutant with sorbitol instead the release of oxygen detected in the absence of this osmolyte may be related to a partial protective effect on other antioxidant systems like catalase as referred above. Concerning to the release of oxygen observed at 10% of glucose either at the exponential phase or during chronological aging, this can be interpreted as the result of enhanced superoxide radical formation, as the superoxide formed due to electron leak from respiration and its reaction with oxygen yields H_2_O_2_, which, in turn, is discomposed into O_2_ and H_2_O by catalase. On this basis, it can be proposed that although causing different longevity, 10% glucose elicited ROS production at similar levels in both WT and *gsh1Δ* cells because of a fully dysfunctional ETC.

*GSH1* mutation affected ROS levels, mitochondrial function, and oxidative stress at 0.5% glucose without having any effect on these parameters at higher glucose concentrations, which suggest that glutathione is essential to preserve mitochondrial function, low ROS production and moderate oxidative stress under dietary restriction. However, it seems that enhanced mitochondrial dysfunction and ROS overproduction caused by low glutathione levels do not have an influence on longevity under dietary restriction since the *GSH1* mutation did not affect the CLS at 0.5% glucose but it did it at 10% glucose. Therefore, it can be hypothesized that other events influenced by glutathione may be responsible for the decrease in CLS at 10% glucose. One probable candidate is the production of excessive quantities of methylglyoxal, a toxic by-product of glycolysis that is highly toxic for cells because of their deleterious effects on DNA and proteins (Vander Jagt *et al.*, 1992). Excessive methylglyoxal is detoxified in part by glyoxalase I, which converts methylglyoxal into S-D-lactoylglutathione in the presence of glutathione (Inoue and Kimura, 1996). Therefore, decreased glutathione synthesis may be disrupting this adaptive pathway and enhancing the deleterious effect of methylglyoxal, which should be expected to be occurring because of the high (i.e. 10%) concentration of glucose in the medium. This is also in agreement with the protection conferred by the exogenous addition of 100 µM glutathione against the deleterious effects of 10% glucose on the CLS of the *gsh1Δ* mutant (Fig 5). Another probable process affecting CLS due to glutathione depletion is the synthesis of iron-sulfur clusters. In this concern, it has been demonstrated that glutathione is important for the maturation of iron sulfur clusters (Kumar *et al.*, 2011). Indeed, the effect of *GSH1* mutation upon iron sulfur proteins may be a cause of the impairment in the mitochondrial respiration observed in this study and the reason for the decrease in the respiratory metabolism in *gsh1Δ* mutant cells grown under dietary restriction (Mannarino *et al.*, 2008). Despite these arguments, it must be taken into account that fermentative metabolism because the culture of yeast in glucose as a carbon source might be buffering the deleterious effects of excessive glucose over mitochondrial function. Thus, the effect of the *GSH1* mutation on CLS and its relationship with mitochondrial dysfunction and oxidative stress deserves further investigation using a non-fermentable carbon source.

Metabolic alterations characterizing metabolic syndrome have been related to glucose-induced oxidative stress (Johnson *et al.*, 2013). Augmented glycolytic flux increases the NADH/NAD^+^ ratio and enhances the reduction of the electron carries from the ETC. This augments mitochondrial membrane potential, which decreases the rate of electron transfer through the ETC leading to electron leak and exacerbated ROS generation. On the contrary, dietary restriction limits the amount of reduced electron carriers and decreases mitochondrial membrane potential, which increases the rate of electron transport to restore the mitochondrial membrane potential, being the final result a more rapid oxygen consumption and lower electron leak and attenuated ROS generation (Murphy, 2009). In agreement with this idea, the data from this study in the WT strain shows that there is an inverse relationship between the concentration of glucose and the rate of mitochondrial respiration, while higher H_2_O_2_ levels were detected, as the concentration of glucose was incremented.

The data from the *gsh1Δ* mutant contradicts the role of mitochondrial dysfunction and excessive ROS generation in the decreased CLS of yeast at 10% glucose. However, the notable increase in H_2_O_2_, the almost full suppression of mitochondrial respiration and the depletion of glutathione at levels comparable to the *gsh1Δ* mutant observed in the WT cells at 10% glucose during the exponential phase (Figs. 1 and 2), might set yeast for a more unfavorable redox environment decreasing its probabilities to survive during chronologic aging, which is in agreement with the notion that in post-diauxic cultures ROS generation occurs inversely to the CLS (Ramos-Gomez *et al.*, 2017).

Despite dietary restriction decreases the levels of oxidative stress in the majority of organisms, it is known that this manipulation has a negligible effect on antioxidant systems, except on glutathione concentration, which remained high in the majority of studies about this issue (Walsh *et al.*, 2014). The data of this report are in agreement with these studies since it was found that glutathione concentration and GSH/GSSG ratio were greater under dietary restriction than in high-glucose level.

Together, these data show that glutathione disturbs CLS in a glucose-dependent manner. Besides, these results provide further insight about the role of glucose in the CLS and its relationship with ROS generation and glutathione redox state.

## Acknowledgments

The authors would like to thank Minerva Ramos-Gomez for their technical support. This work was supported by grants from Instituto Tecnológico Superior de Ciudad Hidalgo (3308.100310) and Tecnológico Nacional de México (165.14.2-PD and 166.14.2-PD).

